# MUTUALISMS ALTER LEGUME NICHES AT A GLOBAL SCALE

**DOI:** 10.64898/2026.05.26.727523

**Authors:** Erin E. McHugh, Megan Bontrager, Megan E. Frederickson

## Abstract

Although mutualism is widely expected to broaden the realized niche of participating species, there have been no global syntheses examining how mutualism affects species’ niches. Here, we combine mutualism trait data on 2,668 legume species with geographic occurrence, climate, and soil nitrogen data. We use these data to investigate the effects of generalized defensive ant-plant mutualisms and more specialized nutritional legume-rhizobium mutualisms on the range of temperature, precipitation, and soil nitrogen values where legumes occur. Legumes engaged in mutualisms with ants or rhizobia occupied significantly more biomes and generally larger niches than non-mutualist species. Legumes with extrafloral nectaries, a trait that attracts a generalized guild of defensive ant partners, had wider niches across all three niche axes, regardless of latitude. Effects of interacting with rhizobia—a more specialized mutualism—on niche breadth were also mainly positive, but depended on latitude, with this mutualism increasing precipitation and soil nitrogen niche breadth only at high latitude, and increasing temperature niche breadth at low latitude, but decreasing temperature niche breadth at high latitude. Mutualism therefore affects niche breadth, but in ways that depend on interaction type and biogeographic trends that vary latitudinally.

## INTRODUCTION

The concept of the niche looms large in ecology. Classically, Hutchinson envisioned a species’ niche as a multidimensional hypervolume in which each axis represents an environmental condition or resource and a species’ traits determine the niche space it can occupy (Hutchinson 1957; Sexton et al. 2017). Hutchinson also distinguished between a species’ fundamental and realized niche, with the latter accounting for species interactions and dispersal limitation (Hutchinson 1957; Futuyma and Moreno 1988; Sexton et al. 2017; Carscadden et al. 2020). The prevailing view of the realized niche has been that it is smaller than the fundamental niche because antagonisms (e.g., competition) limit species to certain habitats or resources they would use otherwise (Hutchinson 1957; Bruno et al. 2003; Sexton et al. 2017; Koffel et al. 2021).

However, mutualisms may expand species’ realized niches beyond their fundamental niches (Bruno et al. 2003; De Mazancourt and Schwartz 2010; Batstone et al. 2018; Koffel et al. 2021). Niche breadth is fundamental to ecology and evolutionary biology, including the study of range limits, range dynamics, and speciation (Sexton et al. 2017). Niche differentiation between lineages can impact speciation (Germain et al. 2021); for example, competition may cause species to evolve niche differences, fueling diversification (Ackerman and Doebeli 2004). Additionally, niche breadth and range size are correlated (Slatyer et al. 2013), and mutualisms may influence range size through direct effects on niche breadth (Afkhami et al. 2014) or because the constricted range of a species geographically restricts the range of its partner (Nuñez et al. 2009; Harrower and Gilbert 2018). The outcome of interplay among mutualism, niche breadth, and range size may be partly driven by the degree of specialization of a mutualism, with generalists more likely to spread to new ranges than specialists (Richardson et al. 2000; Moyano et al. 2020; Fowler et al. 2023; Nathan et al. 2023). Niche breadth plays an outsized role in shaping biodiversity, yet how mutualisms impact niche breadth is still understudied despite the ubiquity of mutualisms in nature (Bruno et al. 2003).

If mutualisms commonly alter realized niche breadth, this would change our understanding of how species interactions shape biogeographic patterns; however, we lack broad empirical tests of this idea. Research to date has been limited to case studies: Afkhami al. (2014) found that engaging with a fungal partner increased the ability of a *Bromus* grass to withstand drought. Kazenel et al. (2015) found that a species of marsh bluegrass that associates with arbuscular mycorrhizal fungi occupies a different niche than a closely related marsh bluegrass with no arbuscular mycorrhizal fungal partner. Forister et al. (2011) found that an ant partner increased survivorship of *Lycaeides melissa* caterpillars on a novel plant host and thus increased *Lycaiedes melissa* host range, likely by reducing predation. These case studies have all found that mutualism increases or shifts niche breadth in specific systems; however, whether this effect is general across taxa, mutualism types, and geographic regions is not known.

This lack of broad-scale investigation makes it difficult to generalize about how mutualism influences niche breadth across different mutualism types and biogeographic regions (but see Luo et al. 2023; Nathan et al. 2023). Some mutualisms are found primarily at certain latitudes (e.g., elaiosomes on ant-dispersed seeds in Mediterranean regions) (Luo et al. 2023), and others are more widely distributed (e.g., animal-mediated pollination) (Richardson et al. 2000). Latitudinal constraints on mutualism may interact with biogeographic patterns that vary across latitude. For example, the increase and decrease in temperature and precipitation variability, respectively, away from the equator (Vázquez and Stevens 2004) may impact not only the benefits and costs of mutualism (Chamberlain et al. 2014), but the magnitude of change in the realized niche of tropical versus temperate species. Additionally, the hypothesized change in stressors from primarily biotic in the tropics to primarily abiotic at high latitudes (Dobzhansky 1950; Louthan et al. 2015), and the often observed increase in the strength of the biotic interactions in the tropics (Schemske et al. 2009, Nathan et al. 2025) may impact how partners benefit their hosts.

Here, we investigate the effects of two well-studied mutualisms on niche breadth in legumes. First, we examine a widespread defensive plant-animal mutualism mediated by extrafloral nectaries (EFNs). EFNs are nectar-secreting plant glands on non-floral tissues that attract ants and some other arthropods, which in turn attack or deter plant herbivores (Bronstein et al. 2006). This interaction is not specialized, with multiple ant species visiting a plant partner (Bronstein et al. 2006; Ness et al. 2009), and is very common both in the tropics (Bronstein et al. 2006) and in legumes (Weber and Keeler 2013). Second, we examine the effects of interacting with rhizobia on legume niche breadth. Rhizobia are nitrogen-fixing bacteria that live in nodules on legume roots, where they exchange fixed nitrogen for fixed carbon from their plant host (Heath and Tiffin 2009). Legume-rhizobium interactions are usually more specialized than legume interactions with ants via EFNs (Heath and Tiffin 2009; Weese et al. 2015; Harrison et al. 2018), making them a useful comparison in this study. Legumes with EFNs have been successfully introduced to more new ranges than legumes lacking EFNs, while specialized legume-rhizobium mutualisms constrain legume introductions (Simonsen et al. 2017; Harrison et al. 2018; Parshuram et al. 2023, Nathan et al. 2023). However, whether these differences in introduction success are due to mutualism-driven changes to legume realized niches is not clear (Fowler et al. 2023; Nathan et al. 2023).

Using global occurrence and climate data, we analyzed three niche variables for mutualist and non-mutualist legumes: precipitation, temperature, and soil nitrogen. Temperature and precipitation are major drivers of biogeographic patterns (Vázquez and Stevens 2004) and can affect mutualistic outcomes. Mutualism strength can be modified by water availability (Pringle et al. 2013) and mutualism may increase or decrease water stress experienced by the host, depending on mutualistic traits (Milligan et al. 2023). Additionally, several studies have found temperature affects mutualism strength, with conflicting results (Barton and Ives 2014; Magnoli et al. 2023). We also characterized the range of soil nitrogen content that species occupy, as rhizobia are most beneficial to legumes under low-nitrogen conditions, and partnerships may evolve to be less cooperative in areas with high soil nitrogen (Heath and Tiffin 2009; Lau et al. 2012; Weese et al. 2015). Additionally, plant-rhizobia interactions may be more common in areas with limited available nitrogen (Vitousek et al. 2013; Sheffer et al. 2015; Menge et al. 2017), which could allow species that engage with rhizobia to inhabit a wider soil nitrogen niche.

We hypothesized that mutualism with rhizobia will lead to decreased precipitation and temperature niche breadths, because legumes engaged in these specialized interactions may be limited in their geographic range or niche size by the availability of compatible, high-quality rhizobia partners (Simonsen et al. 2017; Harrison et al. 2018; Batstone et al. 2018). However, legume-rhizobium mutualisms may increase legume soil nitrogen niche breadth, as receiving nitrogen from rhizobia partners may allow plants to colonize low-nitrogen soils (Sheffer et al. 2015; Menge et al. 2017). In contrast, we predicted that engaging in mutualisms with ants via extrafloral nectaries, a generalized mutualism where plants may have many partners (Bronstein et al. 2006) and often interact with new partners in new areas (Ness et al. 2013; Johnson et al. 2019; Brown and Frederickson 2025), will increase the range of precipitation, temperature, and soil nitrogen ranges that legumes can inhabit. This mutualism provides anti-herbivore defense to plants, and as a result, may reduce biotic stress, broadening species’ abiotic niches (Bronstein et al. 2006; Ness et al. 2009).

## METHODS

### Obtaining data

We began with a legume trait dataset compiled by Simonsen et al. (2017) and used in Harrison et al. (2018) and Nathan et al. (2023). This resource has data on the mutualisms and life-history traits of 3974 legume species. Data on whether species have extrafloral nectaries were originally compiled by Nathan et al. (2023), using the World List of Plants with Extrafloral Nectaries (www.extrafloralnectaries.org). We updated the trait dataset with the latest EFN trait data from this list (Keeler et al. 2026). Data on whether species engage with rhizobia were assembled from Werner et al. (2015) by Simonsen et al. (2017). Simonsen et al. (2017) also used the International Legume Data and Information Service to compile data on whether species are herbaceous or woody, and perennial or annual, as well as how many known human uses (ex., medical, food, or forage uses) they have. However, for both life-history traits, data were not available for every species; thus, missing values were imputed by Simonsen et al. (2017) using characteristics of the species’ genus as indicators of life history. Using *taxize* (Chamberlain et al. 2020), we created a GBIF (GBIF 2024) backbone and filtered the backbone to remove species with fewer than 50 occurrences in GBIF. This left a dataset with 2910 species, for which we downloaded 19,069,248 geographic occurrences from GBIF using *rgbif* (Chamberlain and Boettiger 2017; Chamberlain et al. 2024) on March 2, 2024 (GBIF.org 2024).

### Cleaning data

We used *CoordinateCleaner* (Zizka et al. 2023) to remove occurrences located within capital cities, near or on country centroids, at equal latitude and longitude, near museums, herbaria, and other institutions where cultivated plants and herbarium specimens may be present, at zero degrees latitude and zero degrees longitude, and in the ocean. We used a 1:50m scale to generate *rnaturalearth* maps (Massicotte et al. 2023) to increase, relative to the default, the detail of the maps used to determine which occurrences fell on land versus ocean. Additionally, we included a 25m buffer around all landmasses to account for coastal occurrence points that may have been recorded inaccurately, thus causing them to appear to be in the ocean. These two modifications allowed us to keep more coastal occurrence points in the initial dataset, although they ended up being removed later when extracting terrestrial climate and soil data. To account for observation bias in the data (Inman et al. 2021), we thinned occurrences to one occurrence per square kilometer using *terra* (Hijmans et al. 2024). We used the BioClim 1 raster layer (Fick and Hijmans 2017) (with climate data at a 1 km² resolution) as a guide for thinning. The thinned dataset comprised 6,656,938 occurrences for the 2910 legume species.

### Extracting climate, soil nitrogen, and biome data

We downloaded mean annual temperature and precipitation raster layers (BioClim 1 and BioClim 12) from WorldClim at the resolution of 30 seconds, or 1 km² (Fick and Hijmans 2017). We used *geodata* (Hijmans et al. 2025) to download soil nitrogen data (depth: 5-15 cm, resolution: 250 x 250 m) from SoilGrids, a database that uses predictive modelling to estimate global terrestrial soil nitrogen content (Poggio et al. 2021). Thinned occurrence data was then overlaid with BioClim and SoilGrids (Fig. 1). Fewer than 1% of occurrences lacked corresponding temperature and precipitation values, while 13.4% of occurrences lacked corresponding soil nitrogen values; these missing values were omitted when calculating niche breadth.

**Figure 1.**
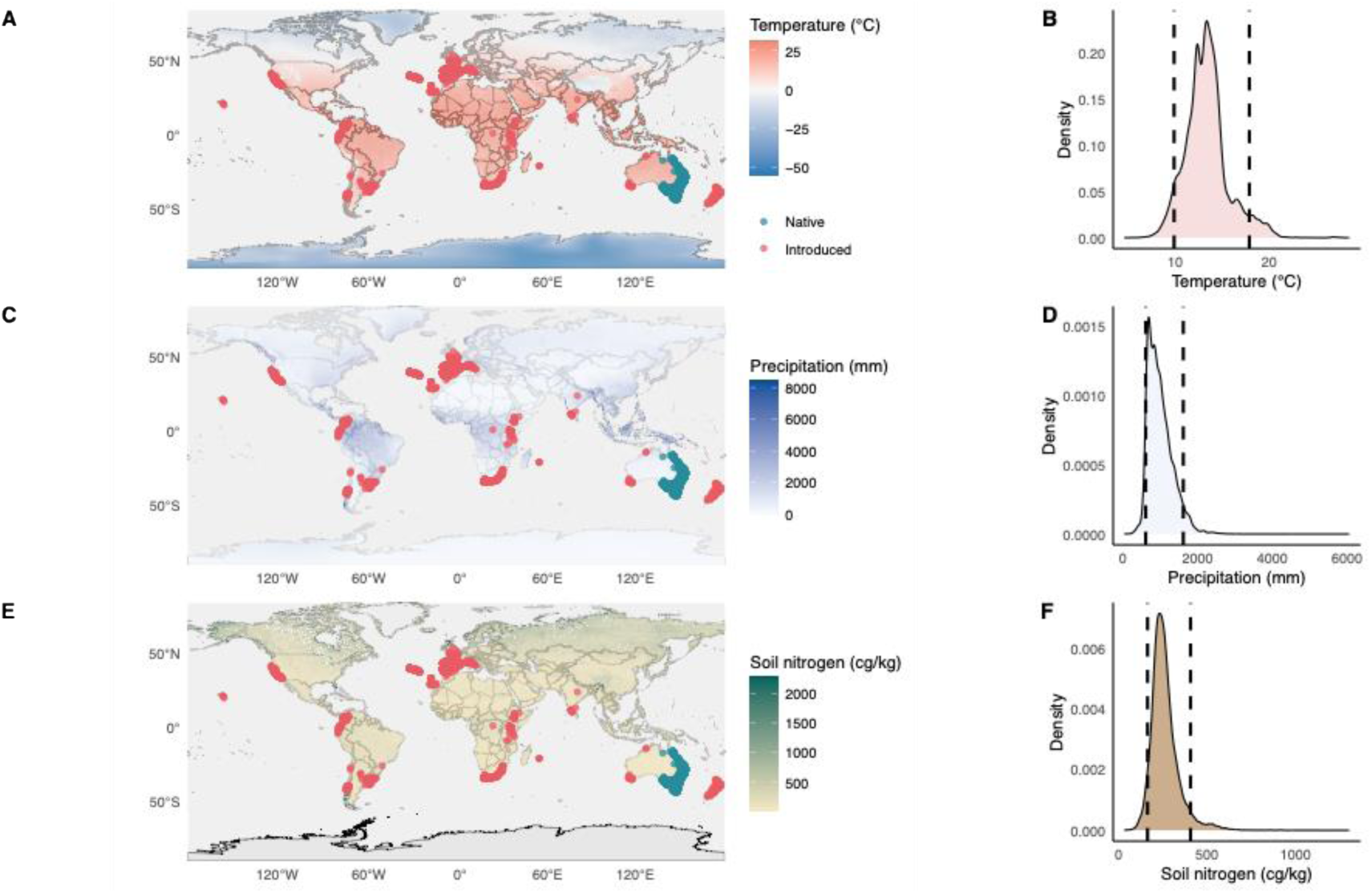
Example of workflow for a single species, *Acacia melanoxylon,* showing native (green dots) and introduced (red dots) GBIF occurrences overlaid on A) mean annual temperature, C) mean annual precipitation, and E) estimated soil nitrogen content raster data to extract values for each occurrence, and the resulting density curves for the distribution of B) mean annual temperature, D) mean annual precipitation, and F) estimated soil nitrogen values where *Acacia melanoxylon* occurs, with dashed lines showing the 5th and 95th percentiles of the distribution.

To obtain the dominant biome for each species, we used WWF’s Terrestrial Ecoregions of the World polygons (Olson et al. 2001), spatially joining occurrences with the biome in which they fall and determining which biome accounted for the largest proportion. From the dataset used to fit our main models, we removed 7,238 and 178 occurrences in the ‘lake’ or ‘rock and ice’ biomes, respectively, as legumes are unlikely to occur in these biomes; this did not omit any species, as no species exclusively occurred in these biomes. We also fit models without filtering occurrences by biome, with very similar results (not shown).

### Reconstructing a phylogenetic tree

To account for phylogenetic signal in the traits of related species (Felsenstein 1985; Symonds and Blomberg 2014), we used *V.phylomaker* (Jin and Qian 2019) to create a phylogenetic tree for all 2910 species in the initial dataset. We used the more conservative nodes.info.1 setting, which creates roots for species clusters only when said cluster is monophyletic (Jin and Qian 2019). A large basal polytomy with 147 tips was dropped using *ape* (Paradis et al. 2024), reducing the number of species in the tree from 2910 to 2763.

### Classifying occurrences as native or introduced

We obtained polygons reflecting the distributions of each species from Plants of the World Online (Govaerts (ed.) 2022). This database uses level three of the World Geographical Scheme for Recording Plant Distributions, in which smaller countries are single polygons, but larger countries are divided into smaller subregions such as provinces and states (Brummitt et al. 2001), each with its own polygon. The database classifies each polygon for each species as either native or introduced. We spatially joined occurrences with their associated polygons (and thus their native or introduced status) with *sf* (Pebesma 2018; Pebesma et al. 2024), using planar geometry. We removed 15 species lacking corresponding spatial polygons from the dataset from our main models, but retained them when modelling unfiltered occurrence data, with similar results (not shown).

A few species in the dataset had poor overlap with the Plants of the World Online polygons, likely due to Plants of the World not capturing the current distribution of the species, or errors in GBIF data. Thus, we filtered the dataset to remove species with fewer than 50% of their occurrence points falling within Plants of the World Online polygons (46 out of 2895 species with Plants of the World Online polygons). We also removed a further 39 species that had fewer than 25 occurrences remaining after cleaning and thinning, reducing the number of species in the dataset to 2810. This filtered dataset was further reduced to 2668 species after removing species not among the 2763 tips in the phylogenetic tree. We also fit models without any of these filtering steps, using data from all 2763 species in the phylogenetic tree for the temperature and precipitation niche breadth models, and from 2762 species for the soil nitrogen niche breadth model, with qualitatively similar results (not shown). One species, a rare island endemic with just four GBIF occurrences (after thinning), did not have any soil nitrogen data.

### Statistical models

First, to understand how mutualism impacts how many biomes a species inhabits, we modelled the number of biomes inhabited by each species as a function of the presence or absence of EFNs, the presence or absence of nodules housing rhizobia, the absolute median latitude of all cleaned and thinned occurrences for each species, and interaction terms between each mutualism state and absolute median latitude. As covariates, we also included plant habit (woody or herbaceous), life history (annual or perennial), and number of human uses of each plant species. We fit phylogenetic generalized least squares models (PGLS), thereby accounting for how phylogenetic relationships create non-independence among legumes (Felsenstein 1985). Phylogeny was incorporated by including the tree in the model as a Pagel correlation structure (Symonds and Blomberg 2014), and lambda was calculated using maximum likelihood. Models were run using *nlme* (Pinheiro et al. 2024), predictions extracted using *ggeffects* (Lüdecke et al. 2025), and data visualization completed using the *tidyverse* (Wickham et al. 2019). Analyses were run using R (R Core Team 2024) in RStudio (RStudio Team 2024).

To characterize associations between mutualism and niche breadth, as well as the maximum and minimum values for each niche dimension, we used the 95th and 5th percentiles of each environmental variable as maximum and minimum niche values, respectively, and the 95th minus the 5th percentile as a measure of niche breadth. Using these percentiles, rather than the absolute maximums and minimums, make our niche characterizations less sensitive to outliers from erroneous GBIF localities or transient individuals occupying areas outside the species’ niche. We also fit models using other percentiles (90/10 and 97.5/2.5) with similar results (not shown). We log-transformed niche breadth variables to improve normality, which is assumed by GLS functions (Mundry 2014; Symonds and Blomberg 2014), and modelled niche variables with the same predictors as in the biome number model described above.

To understand whether mutualism has different effects on niche breadth in native versus introduced ranges, we first fit the niche breadth models described above using native range occurrences only, and omitting occurrences from introduced ranges, if any, for each species. We required that species had at least 25 native occurrences to model niche breadth, reducing the filtered dataset size from 2668 to 2649 species with tips in the phylogeny. Next, we took all species with both a native range and an introduced range (309 species with tips in the phylogeny) and summarized the niche breadth for points in each species’ native and introduced ranges separately. We fit the same PGLS models as described above, with the same covariates, for introduced niche breadth, using lambda outputs from niche breadth models run on the total dataset as fixed lambda values in the models. We also fit models on native niche breadth for the reduced dataset with only 309 species to determine whether the reduced statistical power of the 309 species was enough to detect changes in niche breadth.

## RESULTS

After data cleaning, there were 2668 species in the filtered dataset. 2439 of the species engage in one or both mutualisms, while 229 do not participate in either mutualism (Fig. 2). Specifically, 265 species in the dataset have EFNs and 2403 do not, while 2396 species in the dataset form nodules with rhizobia and 272 species do not. The phylogenetic distribution of mutualism states is shown in Fig. S1. The species with the smallest number of occurrences after thinning, *Vigna triphylla*, had 25 occurrences, while the species with the largest number of occurrences after thinning, *Trifolium repens,* had 343,192 GBIF records.

**Figure 2.**
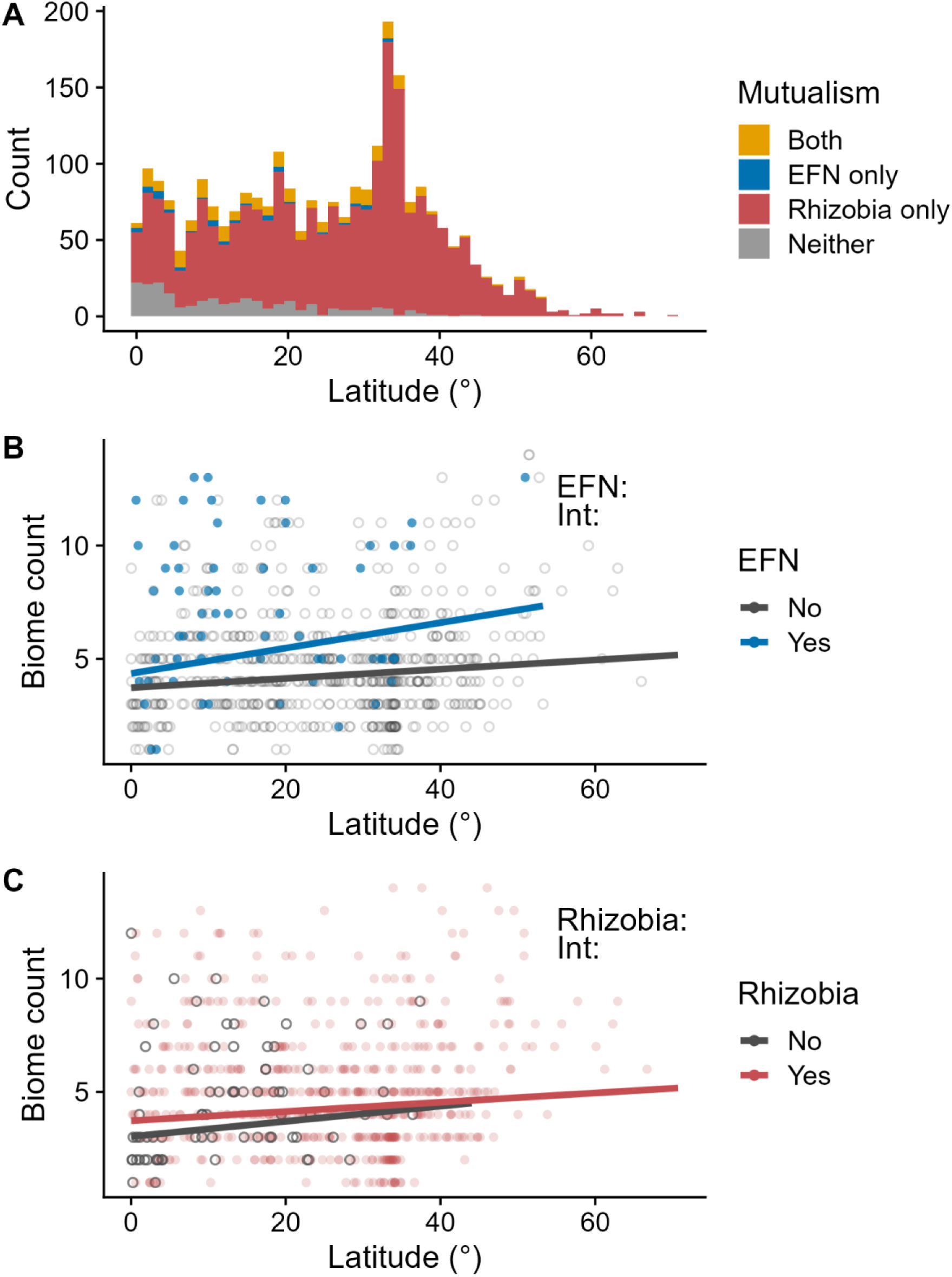
A) Number of legume species with EFNs only (blue), rhizobia only (red), both mutualisms (orange), or neither mutualism (grey) across latitudes. B) PGLS model predictions (lines) plotted over raw data (points) for number of biomes inhabited by species that do (blue) and do not (dark grey) have EFNs or C) do (red) or do not (dark grey) form nodules with rhizobia plotted against the absolute median latitude of the species’ occurrences. To reduce overplotting, only 50% of the raw data points, chosen at random, are shown. Annotations indicate statistical significance of the main effect of EFNs or rhizobia on biome count, or interaction of the mutualism with absolute median latitude (“Int”) (* p < 0.05, ** p < 0.01, *** p < 0.001). See Table S1 for full PGLS model results.

For both mutualism types, mutualist species occupied a significantly larger number of biomes than non-mutualist species (Fig. 2, Table S1). Legumes that make nodules with rhizobia occurred in an estimated 0.696 more biomes than non-nodulating legumes, an effect that did not vary significantly with latitude; there was no significant rhizobia x latitude interaction effect (Fig. 2, Table S1). At the equator, legumes with EFNs occurred in 0.682 more biomes than non-EFN legumes, an effect that intensified with increasing latitude (Fig. 2), indicated by a significantly positive EFN x latitude interaction effect (Table S1). At 30 degrees in latitude, EFN-bearing legumes were predicted to inhabit 1.708 more biomes than legumes without EFNs.

Mutualist species also tended to occupy broader abiotic niches than non-mutualists, although effects varied with latitude for legumes that associate with rhizobia (Fig. 3). Legumes with EFNs had wider niche breadths at all latitudes than legumes without EFNs. Having EFNs was correlated with significantly larger precipitation (Fig. 3A; Table S2), temperature (Fig. 3C; Table S3), and soil nitrogen (Fig. 3E, Table S4) niches, with no significant differences in the effects of this mutualism on niche breadth across latitudes. The wider niche breadth of legumes with EFNs, compared to legumes without EFNs, was due to species with EFNs inhabiting areas with significantly lower minimum annual temperatures than non-EFN species (Fig. S3; Table S9), as well as areas with significantly higher maximum annual precipitation (Fig. S2; Table S5) and soil nitrogen (Fig. S2, Table S7). EFNs did not allow legumes to live in hotter, drier, or more N-poor places; EFNs did not have a significant effect on maximum annual temperature (Table S6), minimum annual precipitation (Table S8), or minimum soil nitrogen (Table S10). As with the effects of EFNs on niche breadth, there were no significant EFN x latitude interactions for the maximum or minimum of any niche variable (Tables S5-S10). In other words, at all latitudes where they occur, legumes with EFNs have wider niches because they grow in cooler, wetter, and more N-rich areas, compared to legumes that lack EFNs.

**Figure 3.**
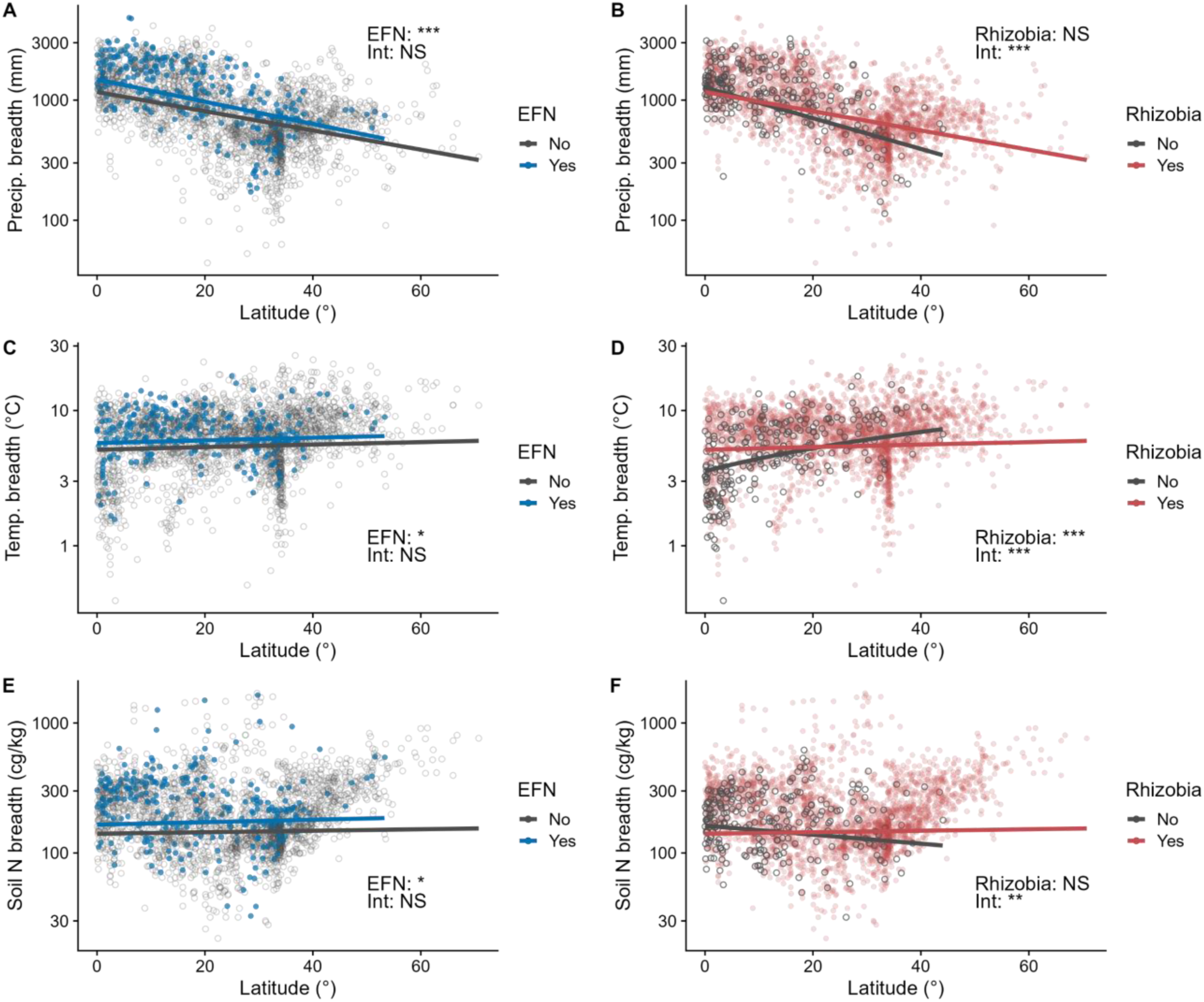
PGLS model predictions (lines) plotted over raw data (points) for precipitation niche breadth (A, B), temperature niche breadth (C, D), and soil nitrogen niche breadth (E, F) against the absolute median latitude of the species’ occurrences, for legume species that do (blue) and do not (dark grey) have EFNs (A, C, E) and do (red) or do not (dark grey) form nodules with rhizobia (B, D, F). Annotations indicate statistical significance of the main effect of EFNs or rhizobia on niche breadth, or interaction of the mutualism with absolute median latitude (“Int”) (* p < 0.05, ** p < 0.01, *** p < 0.001). See Tables S2-S4 for full PGLS model results.

Legumes with symbiotic rhizobia also had different niches compared to non-symbiotic legumes, but effects varied across latitudes. Along precipitation and soil nitrogen niche axes, legumes with rhizobia did not differ from non-symbiotic legumes in niche breadth at the equator but had wider niche breadths at higher latitudes (Fig. 3), indicated by significantly positive rhizobia x latitude interaction terms for precipitation niche breadth (Table S2) and soil nitrogen niche breadth (Table S4). In contrast, legumes with rhizobia had significantly larger temperature niche breadths at the equator, but significantly smaller temperature niche breadths at high latitudes (specifically, above 15.8°) relative to non-symbiotic legumes (Fig. 3, Table S3).

Although rhizobia did not alter legume soil nitrogen and precipitation niche breadth at the equator, shifts in the niche were detected. At the equator, despite no change in soil nitrogen or precipitation niche breadths, legumes with rhizobia occupied drier and more N-poor areas than non-symbiotic legumes (Figs. S2 and S3), indicated by significantly lower 95^th^ and 5^th^ percentiles for both mean annual precipitation (Tables S5 and S8) and soil nitrogen content (Tables S7 and S10). However, this pattern flipped at higher latitudes, with significant rhizobia x latitude interaction terms indicating that legumes with rhizobia grow in significantly wetter and more N-rich areas above ∼13-18° in latitude (Tables S5, S7, S8, and S10). Associating with rhizobia had no main or interactive effect on minimum temperatures inhabited by legume species (Table S9), but legumes with rhizobia inhabited higher maximum temperatures at the equator, but lower maximum temperatures than non-symbiotic legumes at high latitudes (Fig. S3, Table S6). Table S11 has a summary of effects on niche maximums and minimums.

Across all models, absolute median latitude and number of human uses also significantly predicted niche variables. Precipitation (Table S2) and soil nitrogen (Table S4) niche breadth significantly decreased with distance from the equator, changes resulting from decreases in both minimum and maximum precipitation (Tables S5 and S8) and soil nitrogen (Tables S7 and S10) values at high latitudes. In contrast, temperature niche breadth was lowest at the equator and increased at high latitudes (Table S3), a pattern that was still accompanied by decreases in minimum (Table S9) and maximum (Table S6) temperatures with increasing latitude. Number of human uses significantly increased niche breadth across all three axes (Tables S2-S4), with species with more human uses living in both hotter and colder, wetter and drier, and more N-rich and N-poor areas (Tables S5-S10), representing an expansion of the niche. Finally, lambda values ranged from 0.51-0.60 for niche breadth models, 0.57-0.65 for niche maximum models, and 0.64-0.72 for niche minimum models, suggesting relatively strong phylogenetic signal in species’ niche characteristics (Fig. S1).

Niche breadth models using native range occurrences only gave the same results as niche breadth models using all occurrences; all predictors that were significant in models using all occurrences to calculate niche breadth were also significant when calculating niche breadth from native occurrences only (compare Tables S12-S14 to Tables S2-S4). When modelling just the 309 species in the dataset with an introduced range, mutualism was not significantly associated with niche breadth in the introduced range along any axis (Tables S15-S17). However, this analysis likely lacks power to detect effects; models of niche breadth in species’ native ranges using the pared-down dataset with only the same 309 species also found no significant effects of mutualism, although there was a marginally significant main effect of EFNs on soil nitrogen breadth (p=0.0501) (Tables S18-S20). Only 29 of the 309 introduced legume species in the pared-down dataset are non-nodulating, while 62 of the 309 species have EFNs. Compared to non-introduced legumes, these introduced species were more likely to have EFNs (chi-squared: 38.834, p<<0.001), and equally likely to form nodules (chi-squared: 0.160, p= 0.689).

## DISCUSSION

By pairing trait and global occurrence data, we tested the effects of mutualism on legume niche breadth. Legume species with EFNs had wider niche breadth across all three axes. In contrast, legumes that interact with rhizobia had wider temperature niches at the equator, but narrower temperature niches at high latitudes, compared to their non-symbiotic counterparts, and they also had wider precipitation and soil nitrogen niches, but only at high latitudes. Overall, both mutualism types were therefore generally associated with wider niches, except for the narrowing of temperature niche breadth among temperate legumes. Thus, the broadening effect of mutualism on a species’ niche observed in previous case studies (e.g., Forister et al. 2011; Afkhami et al. 2014) and predicted by ecologists (e.g., Bruno et al. 2003) largely holds across a large, diverse plant family at a global scale. Similarly, mutualistic taxa also live in more biomes than non-mutualists (Fig. 2).

Species with wider abiotic niches might be expected to have larger geographic ranges, but our niche breadth results do not perfectly map on to past research on mutualism and range size. Prior work has repeatedly found that, in legumes, generalized mutualisms expand geographic range size, while specialized mutualisms constrain it (Simonsen et al. 2017; Harrison et al. 2018; Parshuram et al. 2022; Delavaux et al. 2022, 2024; Nathan et al. 2023; Brown and Frederickson 2025). Most plants with EFNs recruit diverse ant partners, including novel partners upon arrival in new ranges (Ness 2003; Bronstein et al. 2006; Ness et al. 2009), such that we do not expect a lack of compatible ant partners to limit range expansions of EFN-bearing legumes. Instead, past work has found that plants with EFNs are more likely to be successfully introduced to new regions (Nathan et al. 2023) and to colonize oceanic islands (Brown and Frederickson 2025; Luo et al. 2025) than plants that lack EFNs, and we similarly found that EFN-bearing legumes are over-represented among the introduced species in our dataset, recapitulating past results (Nathan et al. 2023). Our results for the effects of EFNs on legume niche breadth are consistent with this geographic pattern; legumes with EFNs are not constrained by their ant partners in either geographic or niche space. Instead, reaping the benefits of ant protection appears to have allowed EFN-bearing legumes to spread more geographically and along climatic and soil fertility gradients (Johnson et al. 2019; Nathan et al. 2023; Fowler et al. 2023). Although ant diversity and abundance are higher in the tropics (Economo et al. 2018; Schultheiss et al. 2022; Luo et al. 2023), the effect of EFNs on legume niche breadth was constant across latitudes and thus not influenced by the greater diversity and abundance of ant partners nearer the equator.

In contrast, rhizobia are usually more specialized on their host plants than ants, and previous work suggests that legumes that associate with rhizobia are geographically constrained by the availability of compatible symbionts (Simonsen et al. 2017; Delavaux et al. 2022, 2024; Parshuram et al. 2023), especially when the symbiosis is highly specific (Harrison et al. 2018). We might therefore expect legumes with rhizobia to have narrower niches than nonsymbiotic taxa because they are more geographically constrained, but we observed this pattern only for temperature and only for temperate taxa. Unexpectedly, tropical legumes with rhizobia had wider temperature niches, and temperate legumes with rhizobia had wider precipitation and soil N niches than their non-symbiotic counterparts—a pattern inconsistent with the narrower geographic ranges of legumes that engage rhizobia in symbiosis. Observed increases in precipitation and nitrogen niche breadth for temperate legumes with rhizobia may be due to the stresses experienced by legumes at different latitudes: rhizobia primarily reduce abiotic stress (Lau et al. 2012; Weese et al. 2015), but biotic interactions may be a primary stressor near the equator (Dobzhansky 1950; Louthan et al. 2015). Thus, the amelioration of abiotic stress may not necessarily lead to an expansion of the niche in the tropics. At high latitudes, species may be experiencing mainly abiotic stress (Louthan et al. 2015), and so amelioration of abiotic stress by rhizobia may lead to a widening of the abiotic niche. The pattern that we observed for temperature runs counter to this line of reasoning; however, one explanation may be that increased nitrogen availability may make plants more resilient to heat stress (Heckathorn et al. 1996), with species at the equator benefiting more than species at higher, colder latitudes.

We also determined whether changes in niche breadth result from changes to maximum or minimum niche values, or both. Controlling for latitude, legumes with EFNs occur at higher maximum precipitation and soil nitrogen values and lower minimum temperatures than legumes without EFNs and therefore have broader niches. Here and elsewhere, our results do not indicate direction of causation; legumes might evolve EFNs and then expand into new niche space, or lineages with broader niches may be more likely to evolve EFNs. For example, lineages with ranges that extend into wet, nitrogen-rich areas may be better equipped to bear the costs of mutualism (Dutton et al. 2016; Calixto et al. 2021; Luo et al. 2023, but see O’Dowd 1979) or derive more benefits from engaging with ant partners (Bronstein et al. 2006; Louthan et al. 2015), and therefore be more likely to evolve EFNs. Alternatively, already having EFNs may make legumes better able to colonize wet, N-rich habitats despite their high herbivore pressure. The observed pattern of legumes with EFNs inhabiting colder areas than plants without EFNs may be explained by the role that EFNs play in ameliorating biotic stress; when latitude is controlled for, species with EFNs may be better able to inhabit colder areas because protection from herbivory affords them more resources to protect against cold stress.

Tropical legumes that interact with rhizobia were found in areas with lower precipitation, lower nitrogen, and warmer maximum temperatures. At high latitudes, however, the trends were reversed, with species associated with rhizobia found in wetter and more nitrogen-rich areas, and areas with colder maximum temperatures. In the tropics, interacting with rhizobia allowed legumes to live at lower soil nitrogen, as we had hypothesized. As well, nitrogen addition has been shown to decrease the effects of drought on plants (Pivovaroff et al. 2016; Kelso et al. 2020); species that associate with rhizobia have more ready access to plant-available forms of nitrogen, and this nitrogen may confer increased drought resistance, allowing tropical legumes to live in drier regions. Similarly, the increased maximum temperatures observed for tropical legumes that associate with rhizobia may be attributed to the fact that partnering with rhizobia may relieve nutrient stress (Lau et al. 2012; Weese et al. 2015), allowing more resources to tolerate abiotically stressful environments. The reversal of these patterns at higher latitudes is more difficult to explain, although one explanation for why legumes with rhizobia occur in more N-rich soils at some latitudes is that the legume-rhizobium interaction has fertilized the soil; legumes with rhizobia have strong positive effects on soil N cycling and availability (Gou et al. 2023), especially around 28-35°S in Australia and South Africa, where there are many native legumes in shrublands. A spike in species richness of legumes at these latitudes is evident from Fig. 2, suggesting that these legumes may have a large influence on results.

The effects of ant-plant and legume-rhizobium mutualisms on niche breadth were driven by species’ niches in their native ranges. We modelled niche breadth based on both native and introduced occurrences (Tables S2-S4), native occurrences only (Tables S12-S14), and introduced occurrences only (Tables S15-S17), and found that mutualism had the same significant effects in niche breadth models based on both native and introduced occurrences as in models based on native occurrences only. However, mutualism had non-significant effects on niche breadth when considering introduced occurrences only. Nonetheless, it is likely that this smaller dataset (with only 309 introduced species) limited the power to detect differences in niche breadth between mutualistic and non-mutualistic species. Models of niche breadth based on all occurrences of 2668 species or native occurrences only of 2647 species found significant associations between mutualism and niche breadth, but in the pared-down dataset, no associations between mutualism and niche breadth were statistically significant even when modelling native occurrences (Tables S18-S20). Therefore, the lack of significant effects of mutualism on introduced niche breath may be the result of low statistical power.

Mutualism was not the only predictor in our models that was associated with niche breadth. Absolute median latitude was significant in all models, as niche breadth reflected latitudinal trends in temperature, precipitation, and soil N variability in expected directions (Stevens 1989; Saupe et al. 2019; Barandun et al. 2025). Additionally, the number of human uses for a plant species was strongly associated with increased niche breadth, increased maximum niche values, and decreased minimum niche values across all three niche axes. Such findings make sense when human use of plants is viewed as another plant-animal mutualism. For example, Sonoran Desert plants used by Indigenous groups tend to have larger geographic ranges than plants not used by humans (Flower et al. 2021), suggesting that relationships with humans can increase niche breadth. Additionally, species that were woody tended to be found in wetter areas with a wider range of precipitation values, but a narrower range of temperatures, even after controlling for latitude; these trends may reflect the preponderance of woody plants in wet and invariably warm rainforests, compared to other habitat types (e.g., tropical grasslands) at the same latitude. Finally, annual legumes occupied narrower, but hotter temperature niches and more N-poor and drier areas, consistent with their overrepresentation in deserts (Sprent & Gehlot 2010). Thus, mutualism is just one of many traits impacting niche breadth at a global scale.

Of course, global occurrence data have limitations. For example, species occurrences can be biased towards more populous areas, more charismatic species, and wealthier countries and regions (Meyer et al. 2016). Range coverage is important for calculating accurate niche metrics, but difficult to obtain (Meyer et al. 2016). For example, if species occurrences that represent population sinks are included in analyses, our measurements of niche breadth could be wider than they are in reality; conversely, if biases in occurrence data mean that a climatically different portion of the range is not represented in occurrences, our measurements of niche breadth may be narrower than in reality (Lee-Yaw et al. 2022). Consequently, estimates of species niche breadth based on climate and soil nutrient metrics for species’ occurrences may be inaccurate. We thinned occurrences to 1 km^2^ to account for some spatial variation in observation effort, but this still does not fully address biases in the data. However, there is no reason to expect these biases to apply differently to mutualistic versus non-mutualistic plants.

Our results demonstrate that when mutualism impacts niche breadth, it generally works to widen or shift, rather than decrease, breadth, but that the effects of mutualism on niche breadth depend on biogeographical region and mutualism type. Bruno et al. (2003) previously suggested that mutualisms, by reducing stress, could increase the size of the realized niche, perhaps even beyond the fundamental niche, a concept validated by Koffel et al. (2021). Similarly, de Mazancourt and Schwartz (2010) found that mutualism could increase persistence of species in environments with limited resources. This theory had found some empirical support through case studies (Forister et al. 2011; Afkhami et al. 2014; Kazenel et al. 2015) but had yet to be verified by a global synthesis. Over thousands of legume species, we find that engaging with a mutualist partner generally, but not universally, increases the range of environmental conditions where legumes grow, providing global evidence that mutualism can widen niche breadth.

## ACKNOWLEDGEMENTS

The authors thank Marie Josée Fortin, John Stinchcombe, Takuji Usui, Kevin D. Ricks, Damian Hernandez, and Christopher I. Carlson for project and manuscript feedback. E.E.M. was funded by an NSERC Canada Graduate Scholarship - Master’s. Both M.E.F. and M.B acknowledge funding from NSERC Discovery Grants.

## AUTHOR CONTRIBUTIONS

M.E.F. conceived of the initial idea for the project; methods were designed by M.E.F, M. B., and E.E.M. Code for the project was written by E.E.M, with input from M.E.F. and M.B. The manuscript was written by E.E.M., with feedback and revisions from M.E.F. and M.B.

