## Supplemental Materials for "MUTUALISMS ALTER LEGUME NICHES AT A GLOBAL SCALE"

### **Appendix 1**

Supplemental Figures and Tables for: McHugh, E.E., M. Bontrager, and M. E. Frederickson. Mutualisms alter legume niches at a global scale. Under consideration at *Ecology.*

**Supplemental Figure Captions**

**Figure S1.** Phylogenetic tree showing relationships among the 2668 legume species in the filtered dataset, with tip states indicating which species do (blue) and do not (dark grey) have EFNs (innermost ring) and which species do (red) or do not (dark grey) form nodules with rhizobia (second innermost ring). Outer rings plot the species’ precipitation, temperature, and nitrogen niche breadths, centered and scaled to a mean of 0 and a standard deviation of 1.

**Figure S2.** PGLS model predictions (lines) plotted over raw data (points) for the maximum (i.e., 95^th^ percentile) of the precipitation (A, B), temperature (C, D), or soil nitrogen (E, F) distribution against the absolute median latitude of the species’ occurrences, for legume species that do (blue) and do not (dark grey) have EFNs (A, C, E) and do (red) or do not (dark grey) form nodules with rhizobia (B, D, F). Annotations indicate statistical significance of the main effect of EFNs or rhizobia on niche maximums, or interaction of the mutualism with absolute median latitude (“Int”) (* p < 0.05, ** p < 0.01, *** p < 0.001). See Tables S5, S6, and S7 for full model PGLS model results.

**Figure S3.** PGLS model predictions (lines) plotted over raw data (points) for the minimum (i.e., 5^th^ percentile) of the precipitation (A, B), temperature (C, D), or soil nitrogen (E, F) distribution against the absolute median latitude of the species’ occurrences, for legume species that do (blue) and do not (dark grey) have EFNs (A, C, E) and do (red) or do not (dark grey) form nodules with rhizobia (B, D, F). Annotations indicate statistical significance of the main effect of EFNs or rhizobia on niche minimums, or interaction of the mutualism with absolute median latitude (“Int”) (* p < 0.05, ** p < 0.01, *** p < 0.001). See Tables S8, S9, and S10 for full model PGLS model results.

**Figure S1**

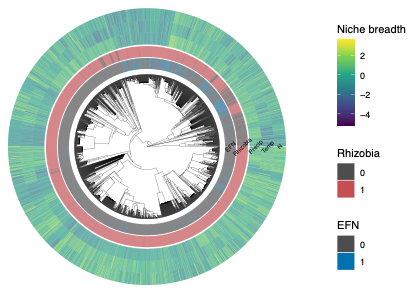

**Figure S2**

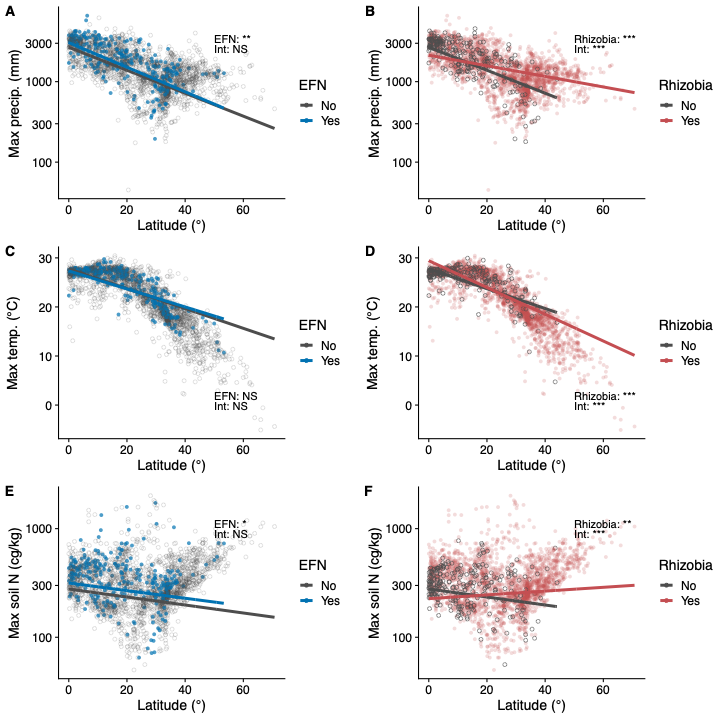

**Figure S3**

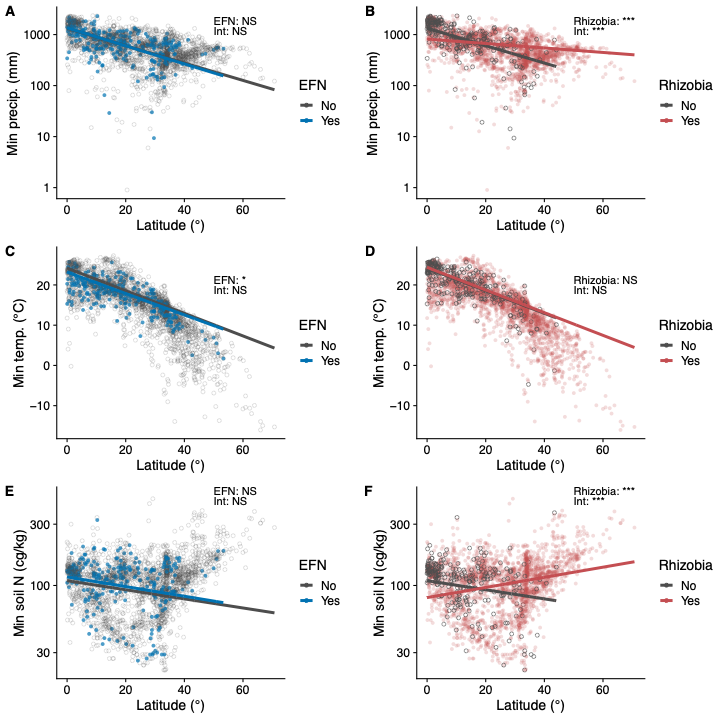

**Table S1.** Effects of mutualism status (EFN no/yes, Rhizobia no/yes), woodiness, life history, number of human uses, and absolute median latitude of species occurrences on biome number. Also included is the interaction between each mutualism status and absolute median latitude. The model intercept represents herbaceous, perennial legumes without EFNs, rhizobia, or any known human uses at the equator. The lambda value estimated by PGLS for this model is λ=0.48. Bold font indicates statistically significant effects.

|  | Value | Standard error | t-value | p-value |
| --- | --- | --- | --- | --- |
| **Intercept** | **3.253** | **0.606** | **5.367** | **0.000** |
| **EFN** | **0.628** | **0.252** | **2.492** | **0.013** |
| **Absolute median latitude** | **0.034** | **0.013** | **2.636** | **0.008** |
| **Rhizobia** | **0.696** | **0.304** | **2.289** | **0.022** |
| **Woody** | **-0.341** | **0.129** | **-2.646** | **0.008** |
| **Number of human uses** | **0.874** | **0.025** | **34.755** | **0.000** |
| Annual | -0.087 | 0.140 | -0.618 | 0.537 |
| **EFN*Absolute median latitude** | **0.036** | **0.011** | **3.371** | **0.001** |
| Rhizobia*Absolute median latitude | -0.013 | 0.013 | -0.999 | 0.318 |

**Table S2.** Effects of mutualism status (EFN no/yes, Rhizobia no/yes), woodiness, life history, number of human uses, and absolute median latitude of species occurrences on log-transformed precipitation niche breadth (95^th^ percentile minus 5^th^ percentile of precipitation distribution). Also included is the interaction between each mutualism status and absolute median latitude. The model intercept represents herbaceous, perennial legumes without EFNs, rhizobia, or any known human uses at the equator. The lambda value estimated by PGLS for this model is λ=0.51. Bold font indicates statistically significant effects.

|  | Value | Standard error | t-value | p-value |
| --- | --- | --- | --- | --- |
| **Intercept** | **7.095** | **0.159** | **44.552** | **0.000** |
| **EFN** | **0.239** | **0.063** | **3.781** | **0.000** |
| **Absolute median latitude** | **-0.029** | **0.003** | **-9.158** | **0.000** |
| Rhizobia | -0.087 | 0.077 | -1.123 | 0.262 |
| **Woody** | **0.103** | **0.032** | **3.173** | **0.002** |
| **Number of human uses** | **0.094** | **0.006** | **14.934** | **0.000** |
| **Annual** | **-0.072** | **0.035** | **-2.062** | **0.039** |
| EFN*Absolute median latitude | -0.003 | 0.003 | -1.077 | 0.282 |
| **Rhizobia*Absolute median latitude** | **0.011** | **0.003** | **3.324** | **0.001** |

**Table S3.** Effects of mutualism status (EFN no/yes, Rhizobia no/yes), woodiness, life history, number of human uses, and absolute median latitude of species occurrences on log-transformed temperature niche breadth (95^th^ percentile minus 5^th^ percentile of temperature distribution). Also included is the interaction between each mutualism status and absolute median latitude. The model intercept represents herbaceous, perennial legumes without EFNs, rhizobia, or any known human uses at the equator. The lambda value estimated by PGLS for this model is λ=0.60. Bold font indicates statistically significant effects.

|  | Value | Standard error | t-value | p-value |
| --- | --- | --- | --- | --- |
| **Intercept** | **1.223** | **0.160** | **7.633** | **0.000** |
| **EFN** | **0.139** | **0.056** | **2.490** | **0.013** |
| **Absolute median latitude** | **0.017** | **0.003** | **5.987** | **0.000** |
| **Rhizobia** | **0.272** | **0.070** | **3.913** | **0.000** |
| Woody | -0.050 | 0.029 | -1.728 | 0.084 |
| **Number of human uses** | **0.060** | **0.006** | **10.896** | **0.000** |
| **Annual** | **-0.091** | **0.031** | **-2.961** | **0.003** |
| EFN*Absolute median latitude | 0.001 | 0.002 | 0.375 | 0.708 |
| **Rhizobia*Absolute median latitude** | **-0.014** | **0.003** | **-4.704** | **0.000** |

**Table S4.** Effects of mutualism status (EFN no/yes, Rhizobia no/yes), woodiness, life history, number of human uses, and absolute median latitude of species occurrences on log-transformed soil nitrogen niche breadth (95^th^ percentile minus 5^th^ percentile of soil nitrogen distribution). Also included is the interaction between each mutualism status and absolute median latitude. The model intercept represents herbaceous, perennial legumes without EFNs, rhizobia, or any known human uses at the equator. The lambda value estimated by PGLS for this model is λ=0.55. Bold font indicates statistically significant effects.

|  | Value | Standard error | t-value | p-value |
| --- | --- | --- | --- | --- |
| **Intercept** | **5.130** | **0.177** | **28.930** | **0.000** |
| **EFN** | **0.162** | **0.067** | **2.425** | **0.015** |
| **Absolute median latitude** | **-0.008** | **0.003** | **-2.338** | **0.020** |
| Rhizobia | -0.143 | 0.082 | -1.738 | 0.082 |
| Woody | 0.001 | 0.034 | 0.020 | 0.984 |
| **Number of human uses** | **0.084** | **0.007** | **12.692** | **0.000** |
| **Annual** | **-0.310** | **0.037** | **-8.386** | **0.000** |
| EFN*Absolute median latitude | 0.001 | 0.003 | 0.303 | 0.762 |
| **Rhizobia*Absolute median latitude** | **0.009** | **0.004** | **2.649** | **0.008** |

**Table S5.** Effects of mutualism status (EFN no/yes, Rhizobia no/yes), woodiness, life history, number of human uses, and absolute median latitude of species occurrences on log-transformed precipitation niche maximum (95^th^ percentile of precipitation distribution). Also included is the interaction between each mutualism status and absolute median latitude. The model intercept represents herbaceous, perennial legumes without EFNs, rhizobia, or any known human uses at the equator. The lambda value estimated by PGLS for this model is λ=0.65. Bold font indicates statistically significant effects.

|  | Value | Standard error | t-value | p-value |
| --- | --- | --- | --- | --- |
| **Intercept** | **7.850** | **0.154** | **50.989** | **0.000** |
| **EFN** | **0.133** | **0.050** | **2.658** | **0.008** |
| **Absolute median latitude** | **-0.033** | **0.003** | **-12.618** | **0.000** |
| **Rhizobia** | **-0.228** | **0.063** | **-3.610** | **0.000** |
| **Woody** | **0.101** | **0.026** | **3.900** | **0.000** |
| **Number of human uses** | **0.051** | **0.005** | **10.381** | **0.000** |
| **Annual** | **-0.155** | **0.027** | **-5.629** | **0.000** |
| EFN*Absolute median latitude | -0.002 | 0.002 | -1.109 | 0.268 |
| **Rhizobia*Absolute median latitude** | **0.018** | **0.003** | **6.631** | **0.000** |

**Table S6.** Effects of mutualism status (EFN no/yes, Rhizobia no/yes), woodiness, life history, number of human uses, and absolute median latitude of species occurrences on log-transformed temperature niche maximum (95^th^ percentile of temperature distribution). Also included is the interaction between each mutualism status and absolute median latitude. The model intercept represents herbaceous, perennial legumes without EFNs, rhizobia, or any known human uses at the equator. The lambda value estimated by PGLS for this model is λ=0.57. Bold font indicates statistically significant effects.

|  | Value | Standard error | t-value | p-value |
| --- | --- | --- | --- | --- |
| **Intercept** | **27.334** | **0.850** | **32.161** | **0.000** |
| EFN | -0.402 | 0.312 | -1.289 | 0.198 |
| **Absolute median latitude** | **-0.202** | **0.016** | **-12.629** | **0.000** |
| **Rhizobia** | **1.668** | **0.386** | **4.325** | **0.000** |
| Woody | 0.150 | 0.160 | 0.933 | 0.351 |
| **Number of human uses** | **0.284** | **0.031** | **9.147** | **0.000** |
| **Annual** | **2.386** | **0.172** | **13.842** | **0.000** |
| EFN*Absolute median latitude | 0.018 | 0.013 | 1.387 | 0.165 |
| **Rhizobia*Absolute median latitude** | **-0.071** | **0.016** | **-4.313** | **0.000** |

**Table S7.** Effects of mutualism status (EFN no/yes, Rhizobia no/yes), woodiness, life history, number of human uses, and absolute median latitude of species occurrences on log-transformed soil nitrogen niche maximum (95^th^ percentile of soil nitrogen distribution). Also included is the interaction between each mutualism status and absolute median latitude. The model intercept represents herbaceous, perennial legumes without EFNs, rhizobia, or any known human uses at the equator. The lambda value estimated by PGLS for this model is λ=0.63. Bold font indicates statistically significant effects.

|  | Value | Standard error | t-value | p-value |
| --- | --- | --- | --- | --- |
| **Intercept** | **5.632** | **0.163** | **34.658** | **0.000** |
| **EFN** | **0.127** | **0.054** | **2.343** | **0.019** |
| **Absolute median latitude** | **-0.008** | **0.003** | **-3.006** | **0.003** |
| **Rhizobia** | **-0.205** | **0.068** | **-2.996** | **0.003** |
| **Woody** | **0.055** | **0.028** | **1.964** | **0.050** |
| **Number of human uses** | **0.052** | **0.005** | **9.568** | **0.000** |
| **Annual** | **-0.289** | **0.030** | **-9.691** | **0.000** |
| EFN*Absolute median latitude | 0.000 | 0.002 | 0.179 | 0.858 |
| **Rhizobia*Absolute median latitude** | **0.012** | **0.003** | **4.312** | **0.000** |

**Table S8.** Effects of mutualism status (EFN no/yes, Rhizobia no/yes), woodiness, life history, number of human uses, and absolute median latitude of species occurrences on log-transformed precipitation niche minimum (5^th^ percentile of precipitation distribution). Also included is the interaction between each mutualism status and absolute median latitude. The model intercept represents herbaceous, perennial legumes without EFNs, rhizobia, or any known human uses at the equator. The lambda value estimated by PGLS for this model is λ=0.65. Bold font indicates statistically significant effects.

|  | Value | Standard error | t-value | p-value |
| --- | --- | --- | --- | --- |
| **Intercept** | **7.174** | **0.228** | **31.437** | **0.000** |
| EFN | 0.021 | 0.075 | 0.276 | 0.783 |
| **Absolute median latitude** | **-0.039** | **0.004** | **-10.081** | **0.000** |
| **Rhizobia** | **-0.484** | **0.094** | **-5.126** | **0.000** |
| **Woody** | **0.094** | **0.039** | **2.441** | **0.015** |
| **Number of human uses** | **-0.017** | **0.007** | **-2.309** | **0.021** |
| **Annual** | **-0.311** | **0.041** | **-7.568** | **0.000** |
| EFN*Absolute median latitude | -0.001 | 0.003 | -0.481 | 0.631 |
| **Rhizobia*Absolute median latitude** | **0.029** | **0.004** | **7.285** | **0.000** |

**Table S9.** Effects of mutualism status (EFN no/yes, Rhizobia no/yes), woodiness, life history, number of human uses, and absolute median latitude of species occurrences on log-transformed temperature niche minimum (5^th^ percentile of temperature distribution). Also included is the interaction between each mutualism status and absolute median latitude. The model intercept represents herbaceous, perennial legumes without EFNs, rhizobia, or any known human uses at the equator. The lambda value estimated by PGLS for this model is λ=0.64. Bold font indicates statistically significant effects.

|  | Value | Standard error | t-value | p-value |
| --- | --- | --- | --- | --- |
| **Intercept** | **23.456** | **1.326** | **17.693** | **0.000** |
| **EFN** | **-0.990** | **0.437** | **-2.266** | **0.024** |
| **Absolute median latitude** | **-0.281** | **0.023** | **-12.354** | **0.000** |
| Rhizobia | 0.184 | 0.552 | 0.334 | 0.738 |
| **Woody** | **0.493** | **0.226** | **2.179** | **0.029** |
| **Number of human uses** | **-0.141** | **0.043** | **-3.257** | **0.001** |
| **Annual** | **3.100** | **0.241** | **12.887** | **0.000** |
| EFN*Absolute median latitude | 0.015 | 0.018 | 0.814 | 0.416 |
| Rhizobia*Absolute median latitude | -0.001 | 0.023 | -0.026 | 0.980 |

**Table S10.** Effects of mutualism status (EFN no/yes, Rhizobia no/yes), woodiness, life history, number of human uses, and absolute median latitude of species occurrences on log-transformed soil nitrogen niche minimum (5^th^ percentile of soil nitrogen distribution). Also included is the interaction between each mutualism status and absolute median latitude. The model intercept represents herbaceous, perennial legumes without EFNs, rhizobia, or any known human uses at the equator. The lambda value estimated by PGLS for this model is λ=0.72. Bold font indicates statistically significant effects.

|  | Value | Standard error | t-value | p-value |
| --- | --- | --- | --- | --- |
| **Intercept** | **4.631** | **0.175** | **26.431** | **0.000** |
| EFN | 0.067 | 0.051 | 1.305 | 0.192 |
| **Absolute median latitude** | **-0.008** | **0.003** | **-3.005** | **0.003** |
| **Rhizobia** | **-0.304** | **0.066** | **-4.590** | **0.000** |
| **Woody** | **0.148** | **0.027** | **5.561** | **0.000** |
| **Number of human uses** | **-0.019** | **0.005** | **-3.650** | **0.000** |
| **Annual** | **-0.238** | **0.028** | **-8.492** | **0.000** |
| EFN*Absolute median latitude | 0.000 | 0.002 | -0.229 | 0.819 |
| **Rhizobia*Absolute median latitude** | **0.017** | **0.003** | **6.195** | **0.000** |

**Table S11.** Summary of how mutualisms impact the breadth, minimum, and maximum of a given niche axis in phylogenetic generalized least squares models.

| Mutualism | Axis | Niche breadth | | Niche maximum | | Niche minimum | |
| --- | --- | --- | --- | --- | --- | --- | --- |
|  |  | Equator | High latitude | Equator | High latitude | Equator | High latitude |
| EFN | Precipitation | + | + | + | + | NS | NS |
|  | Temperature | + | + | NS | NS | - | - |
|  | Soil nitrogen | + | + | + | + | NS | NS |
| Rhizobia | Precipitation | NS | + | - | + | - | + |
|  | Temperature | + | - | + | - | NS | NS |
|  | Soil nitrogen | NS | + | - | + | - | + |

**Table S12.** Effects of mutualism status (EFN no/yes, Rhizobia no/yes), woodiness, life history, number of human uses, and absolute median latitude of species occurrences on log-transformed precipitation niche breadth (95^th^ percentile minus 5^th^ percentile of precipitation distribution) calculated from occurrences in each species’ native range only. Also included is the interaction between each mutualism status and absolute median latitude. The model intercept represents herbaceous, perennial legumes without EFNs, rhizobia, or any known human uses at the equator. The lambda value estimated by PGLS for this model is λ=0.53. Bold font indicates statistically significant effects.

|  | Value | Standard error | t-value | p-value |
| --- | --- | --- | --- | --- |
| **Intercept** | **7.057** | **0.172** | **41.140** | **0.000** |
| **EFN** | **0.294** | **0.067** | **4.410** | **0.000** |
| **Absolute median latitude** | **-0.030** | **0.003** | **-8.836** | **0.000** |
| Rhizobia | -0.072 | 0.082 | -0.885 | 0.376 |
| **Woody** | **0.099** | **0.034** | **2.903** | **0.004** |
| **Number of human uses** | **0.087** | **0.007** | **13.051** | **0.000** |
| **Annual** | **-0.091** | **0.037** | **-2.478** | **0.013** |
| EFN*Absolute median latitude | -0.004 | 0.003 | -1.589 | 0.112 |
| **Rhizobia*Absolute median latitude** | **0.012** | **0.003** | **3.414** | **0.001** |

**Table S13.** Effects of mutualism status (EFN no/yes, Rhizobia no/yes), woodiness, life history, number of human uses, and absolute median latitude of species occurrences on log-transformed temperature niche breadth (95^th^ percentile minus 5^th^ percentile of temperature distribution) calculated from occurrences in each species’ native range only. Also included is the interaction between each mutualism status and absolute median latitude. The model intercept represents herbaceous, perennial legumes without EFNs, rhizobia, or any known human uses at the equator. The lambda value estimated by PGLS for this model is λ=0.62. Bold font indicates statistically significant effects.

|  | Value | Standard error | t-value | p-value |
| --- | --- | --- | --- | --- |
| **Intercept** | **1.173** | **0.167** | **7.037** | **0.000** |
| **EFN** | **0.159** | **0.057** | **2.761** | **0.006** |
| **Absolute median latitude** | **0.018** | **0.003** | **6.042** | **0.000** |
| **Rhizobia** | **0.296** | **0.072** | **4.107** | **0.000** |
| Woody | -0.050 | 0.029 | -1.698 | 0.090 |
| **Number of human uses** | **0.049** | **0.006** | **8.507** | **0.000** |
| **Annual** | **-0.096** | **0.032** | **-3.054** | **0.002** |
| EFN*Absolute median latitude | 0.001 | 0.002 | 0.559 | 0.576 |
| **Rhizobia*Absolute median latitude** | **-0.015** | **0.003** | **-4.925** | **0.000** |

**Table S14.** Effects of mutualism status (EFN no/yes, Rhizobia no/yes), woodiness, life history, number of human uses, and absolute median latitude of species occurrences on log-transformed soil nitrogen niche breadth (95^th^ percentile minus 5^th^ percentile of soil nitrogen distribution) calculated from occurrences in each species’ native range only. Also included is the interaction between each mutualism status and absolute median latitude. The model intercept represents herbaceous, perennial legumes without EFNs, rhizobia, or any known human uses at the equator. The lambda value estimated by PGLS for this model is λ=0.55. Bold font indicates statistically significant effects.

|  | Value | Standard error | t-value | p-value |
| --- | --- | --- | --- | --- |
| **Intercept** | **5.102** | **0.180** | **28.296** | **0.000** |
| **EFN** | **0.142** | **0.068** | **2.087** | **0.037** |
| **Absolute median latitude** | **-0.008** | **0.003** | **-2.220** | **0.027** |
| Rhizobia | -0.160 | 0.084 | -1.907 | 0.057 |
| Woody | 0.018 | 0.035 | 0.509 | 0.611 |
| **Number of human uses** | **0.066** | **0.007** | **9.806** | **0.000** |
| **Annual** | **-0.299** | **0.037** | **-7.969** | **0.000** |
| EFN*Absolute median latitude | 0.002 | 0.003 | 0.559 | 0.576 |
| **Rhizobia*Absolute median latitude** | **0.009** | **0.004** | **2.576** | **0.010** |

**Table S15.** For 309 introduced species only, effects of mutualism status (EFN no/yes, Rhizobia no/yes), woodiness, life history, number of human uses, and absolute median latitude of species occurrences on log-transformed precipitation niche breadth (95^th^ percentile minus 5^th^ percentile of precipitation distribution) calculated from occurrences in each species’ introduced range only. Also included is the interaction between each mutualism status and absolute median latitude. The model intercept represents herbaceous, perennial legumes without EFNs, rhizobia, or any known human uses at the equator. Bold font indicates statistically significant effects.

|  | Value | Standard error | t-value | p-value |
| --- | --- | --- | --- | --- |
| **Intercept** | 7.461 | 0.279 | 26.744 | 0.000 |
| EFN | 0.054 | 0.154 | 0.348 | 0.728 |
| Absolute median latitude | -0.027 | 0.008 | -3.289 | 0.001 |
| Rhizobia | -0.016 | 0.246 | -0.066 | 0.948 |
| Woody | 0.046 | 0.089 | 0.514 | 0.608 |
| **Number of human uses** | **0.038** | **0.013** | **2.916** | **0.004** |
| Annual | -0.035 | 0.084 | -0.420 | 0.675 |
| EFN*Absolute median latitude | 0.001 | 0.005 | 0.241 | 0.810 |
| Rhizobia*Absolute median latitude | 0.003 | 0.008 | 0.315 | 0.753 |

**Table S16.** For 309 introduced species only, effects of mutualism status (EFN no/yes, Rhizobia no/yes), woodiness, life history, number of human uses, and absolute median latitude of species occurrences on log-transformed temperature niche breadth (95^th^ percentile minus 5^th^ percentile of temperature distribution) calculated from occurrences in each species’ introduced range only. Also included is the interaction between each mutualism status and absolute median latitude. The model intercept represents herbaceous, perennial legumes without EFNs, rhizobia, or any known human uses at the equator. Bold font indicates statistically significant effects.

|  | Value | Standard error | t-value | p-value |
| --- | --- | --- | --- | --- |
| **Intercept** | **1.753** | **0.274** | **6.399** | **0.000** |
| EFN | -0.001 | 0.142 | -0.005 | 0.996 |
| Absolute median latitude | -0.009 | 0.008 | -1.125 | 0.261 |
| Rhizobia | 0.022 | 0.234 | 0.093 | 0.926 |
| Woody | -0.008 | 0.082 | -0.098 | 0.922 |
| **Number of human uses** | **0.062** | **0.012** | **5.228** | **0.000** |
| Annual | -0.052 | 0.076 | -0.676 | 0.500 |
| EFN*Absolute median latitude | 0.002 | 0.005 | 0.373 | 0.710 |
| Rhizobia*Absolute median latitude | 0.003 | 0.008 | 0.424 | 0.672 |

**Table S17.** For 309 introduced species only, effects of mutualism status (EFN no/yes, Rhizobia no/yes), woodiness, life history, number of human uses, and absolute median latitude of species occurrences on log-transformed soil nitrogen niche breadth (95^th^ percentile minus 5^th^ percentile of soil nitrogen distribution) calculated from occurrences in each species’ introduced range only. Also included is the interaction between each mutualism status and absolute median latitude. The model intercept represents herbaceous, perennial legumes without EFNs, rhizobia, or any known human uses at the equator. Bold font indicates statistically significant effects.

|  | Value | Standard error | t-value | p-value |
| --- | --- | --- | --- | --- |
| **Intercept** | **5.410** | **0.290** | **18.670** | **0.000** |
| EFN | 0.267 | 0.156 | 1.713 | 0.088 |
| Absolute median latitude | -0.002 | 0.008 | -0.199 | 0.842 |
| Rhizobia | -0.048 | 0.252 | -0.189 | 0.850 |
| Woody | 0.017 | 0.090 | 0.189 | 0.850 |
| Number of human uses | 0.022 | 0.013 | 1.668 | 0.096 |
| **Annual** | **-0.220** | **0.084** | **-2.612** | **0.010** |
| EFN*Absolute median latitude | -0.007 | 0.005 | -1.424 | 0.156 |
| Rhizobia*Absolute median latitude | 0.008 | 0.009 | 0.908 | 0.365 |

**Table S18.** For 309 introduced species only, effects of mutualism status (EFN no/yes, Rhizobia no/yes), woodiness, life history, number of human uses, and absolute median latitude of species occurrences on log-transformed precipitation niche breadth (95^th^ percentile minus 5^th^ percentile of precipitation distribution) calculated from occurrences in each species’ native range only. Also included is the interaction between each mutualism status and absolute median latitude. The model intercept represents herbaceous, perennial legumes without EFNs, rhizobia, or any known human uses at the equator. Bold font indicates statistically significant effects.

|  | Value | Standard error | t-value | p-value |
| --- | --- | --- | --- | --- |
| **Intercept** | **7.318** | **0.236** | **30.963** | **0.000** |
| EFN | 0.200 | 0.127 | 1.573 | 0.117 |
| **Absolute median latitude** | **-0.025** | **0.008** | **-3.270** | **0.001** |
| Rhizobia | 0.124 | 0.206 | 0.601 | 0.548 |
| Woody | -0.041 | 0.070 | -0.582 | 0.561 |
| **Number of human uses** | **0.043** | **0.010** | **4.234** | **0.000** |
| Annual | -0.114 | 0.066 | -1.720 | 0.087 |
| EFN*Absolute median latitude | -0.007 | 0.004 | -1.722 | 0.086 |
| Rhizobia*Absolute median latitude | 0.007 | 0.008 | 0.907 | 0.365 |

**Table S19.** For 309 introduced species only, effects of mutualism status (EFN no/yes, Rhizobia no/yes), woodiness, life history, number of human uses, and absolute median latitude of species occurrences on log-transformed temperature niche breadth (95^th^ percentile minus 5^th^ percentile of temperature distribution) calculated from occurrences in each species’ native range only. Also included is the interaction between each mutualism status and absolute median latitude. The model intercept represents herbaceous, perennial legumes without EFNs, rhizobia, or any known human uses at the equator. Bold font indicates statistically significant effects.

|  | Value | Standard error | t-value | p-value |
| --- | --- | --- | --- | --- |
| **Intercept** | **1.884** | **0.204** | **9.258** | **0.000** |
| EFN | 0.137 | 0.104 | 1.314 | 0.190 |
| Absolute median latitude | 0.001 | 0.006 | 0.134 | 0.894 |
| Rhizobia | 0.192 | 0.172 | 1.116 | 0.265 |
| Woody | -0.112 | 0.058 | -1.942 | 0.053 |
| **Number of human uses** | **0.023** | **0.008** | **2.844** | **0.005** |
| Annual | -0.012 | 0.053 | -0.222 | 0.825 |
| EFN*Absolute median latitude | -0.001 | 0.003 | -0.296 | 0.768 |
| Rhizobia*Absolute median latitude | -0.002 | 0.006 | -0.327 | 0.744 |

**Table S20.** For 309 introduced species only, effects of mutualism status (EFN no/yes, Rhizobia no/yes), woodiness, life history, number of human uses, and absolute median latitude of species occurrences on log-transformed soil nitrogen niche breadth (95^th^ percentile minus 5^th^ percentile of soil nitrogen distribution) calculated from occurrences in each species’ native range only. Also included is the interaction between each mutualism status and absolute median latitude. The model intercept represents herbaceous, perennial legumes without EFNs, rhizobia, or any known human uses at the equator. Bold font indicates statistically significant effects.

|  | Value | Standard error | t-value | p-value |
| --- | --- | --- | --- | --- |
| **Intercept** | **5.477** | **0.263** | **20.856** | **0.000** |
| **EFN** | **0.272** | **0.138** | **1.967** | **0.050** |
| Absolute median latitude | -0.005 | 0.008 | -0.629 | 0.530 |
| Rhizobia | -0.058 | 0.226 | -0.255 | 0.799 |
| Woody | -0.140 | 0.077 | -1.832 | 0.068 |
| **Number of human uses** | **0.024** | **0.011** | **2.183** | **0.030** |
| **Annual** | **-0.318** | **0.071** | **-4.473** | **0.000** |
| EFN*Absolute median latitude | -0.004 | 0.005 | -0.891 | 0.373 |
| Rhizobia*Absolute median latitude | 0.014 | 0.009 | 1.606 | 0.109 |
